# Schizophrenia risk from locus-specific human endogenous retroviruses

**DOI:** 10.1101/798017

**Authors:** Rodrigo R.R. Duarte, Matthew L. Bendall, Miguel de Mulder, Christopher E. Ormsby, Greta A. Beckerle, Sashika Selvackadunco, Claire Troakes, Gustavo Reyes-Terán, Keith A. Crandall, Deepak P. Srivastava, Douglas F. Nixon, Timothy R. Powell

**Affiliations:** Social, Genetic & Developmental Psychiatry Centre, Institute of Psychiatry, Psychology & Neuroscience, King’s College London, London, UK; Department of Basic & Clinical Neuroscience, Institute of Psychiatry, Psychology & Neuroscience, King’s College London, London, UK; Division of Infectious Diseases, Weill Cornell Medicine, Cornell University, New York, NY, USA; Computational Biology Institute, Milken Institute School of Public Health, The George Washington University, Washington, DC, USA; Center for Research in Infectious Diseases, National Institute of Respiratory Diseases, Mexico City, Mexico; MRC London Neurodegenerative Diseases Brain Bank, Institute of Psychiatry, Psychology & Neuroscience, King’s College London, London, United Kingdom; MRC Centre for Neurodevelopmental Disorders, King’s College London, London, United Kingdom

## Abstract

Schizophrenia genome-wide association studies highlight the substantial contribution of risk attributed to the non-coding genome where human endogenous retroviruses (HERVs) are encoded. These ancient viral elements have previously been overlooked in genetic and transcriptomic studies due to their poor annotation and repetitive nature. Using a new, comprehensive HERV annotation, we found that the fraction of the genome where HERVs are located (the ‘retrogenome’) is enriched for schizophrenia risk variants, and that there are 148 disparate HERVs involved in susceptibility. Analysis of RNA-sequencing data from the dorsolateral prefrontal cortex of 259 schizophrenia cases and 279 controls from the CommonMind Consortium showed that HERVs are actively expressed in the brain (n = 3,979), regulated in *cis* by common genetic variants (n = 1,759), and differentially expressed in patients (n = 81). Convergent analyses implicate LTR25_6q21 and ERVLE_8q24.3h as HERVs of etiological relevance to schizophrenia, which are co-regulated with genes involved in neuronal and mitochondrial function, respectively. Our findings provide a strong rationale for exploring the retrogenome and the expression of these locus-specific HERVs as novel risk factors for schizophrenia and potential diagnostic biomarkers and treatment targets.

## Introduction

Human endogenous retroviruses (HERVs) are remnants of genetic material acquired through our evolutionary past which originated from the infection of germline cells with ancient retroviruses. These viruses multiplied through a copy-and-paste mechanism and were eventually endogenized (i.e., vertically transmitted), and now constitute approximately 8% of the genome^1,2^. These repetitive sequences were generally assumed to be transcriptionally inactive in the modern genome, having a purely regulatory function due to the retainment of the viral promoters (long-terminal repeats, LTRs). However, certain HERVs were co-opted to serve novel specialized roles, including in the regulation of embryonic development^3,4^ and neural progenitor cells^5,6^. They have also been implicated in neuropsychiatric conditions such as amyotrophic lateral sclerosis^7–9^, major depressive disorder, bipolar disorder, and schizophrenia^10–12^. Despite their abundance in the genome and relevance to disease and fundamental aspects of human biology, the location and function of most HERVs remain elusive to-date.

Recent large genome-wide association studies (GWAS) comparing schizophrenia cases and non-affected individuals enabled the identification of polymorphisms mediating risk for this disorder^13–15^. These studies highlight the substantial contribution of risk attributed to the non-coding genome, where HERVs are encoded, which are often overlooked in both genomic and transcriptomic studies^16^. Until recently, there was no comprehensively annotated map of HERVs in the genome, and there were no computationally efficient tools to analyze the expression of these repetitive sequences with single-locus resolution. Consequently, previous studies were unable to test whether locus-specific HERVs were genetically associated with traits of interest, or to assess their expression in conditions of interest, or to distinguish expressed from dormant HERVs from within the same family, which contributed to the generation of inconclusive findings. For example, Karlsson and colleagues^17^ reported decreased ERV9 expression in the brain of schizophrenia patients, whereas Diem and collaborators^18^ found the opposite. While the heterogeneity of schizophrenia is also likely to be a contributing factor for these contradictory findings, there are approximately 120 copies of ERV9 in the genome^19,20^ that likely also confounded these reports.

Recently, there have been significant advances in the annotation of HERVs in the genome^20–24^, and in the development of tools that are able to map repetitive sequences in RNA-sequencing data to their most likely source of origin in the genome^20,25,26^. These improvements allow for the identification and quantification of specific HERV copies associated with traits of interest. These advances, combined with modern population genetic methods, enabled us to identify the contribution of the ‘retrogenome’ (the full genomic complement of HERVs) and expression of locus-specific HERVs to schizophrenia etiology and neurobiology.

## Results

### The contribution of polymorphisms in the retrogenome to schizophrenia risk

A recently developed annotation of putatively functional HERV elements describes 14,968 elements dispersed throughout the genome, which were defined based on the presence of LTRs and remnants of genetic elements which code for viral proteins like *env*, *gag* or *pol*^20^. We assessed the contribution of common genetic variants within HERVs to schizophrenia susceptibility using GARFIELD^27^. This pipeline quantifies the co-localization of GWAS results with variants in annotation categories (i.e. variants in the retrogenome), assessing significance using linear models that control for minor allele frequency and linkage disequilibrium. For comparison, enrichment was also calculated for heritable neuropsychiatric traits and non-neuropsychiatric phenotypes: height, body mass index, coronary artery disease, Crohn’s Disease, type 2 diabetes, neuroticism, eczema, major depressive disorder, bipolar disorder, Alzheimer’s disease, attention deficit hyperactivity disorder, amyotrophic lateral sclerosis, and autism spectrum disorder (**Figure 1**; full results on **Supplemental Table 1**). Polymorphisms within HERVs were significantly enriched for genome-wide significant variants (P < 5.00 × 10^−8^) associated with schizophrenia (P_enrichment_ = 1.88 × 10^−5^, P_corrected_ [for 14 traits and 2 GWAS thresholds tested] = 5.27 × 10^−4^, β = 0.90, 95% CI [0.49, 1.31]). Analysis of variants associated with these traits under a more relaxed P cut-off (P < 5.00 × 10^−5^) also showed an enrichment for schizophrenia only (P_enrichment_ = 4.56 × 10^−7^, P_corrected_ = 1.28 × 10^−5^, β = 0.57, 95% CI [0.35, 0.80]). These findings suggest a role for the retrogenome in the etiology of this neurodevelopmental disorder.

**Figure 1.**
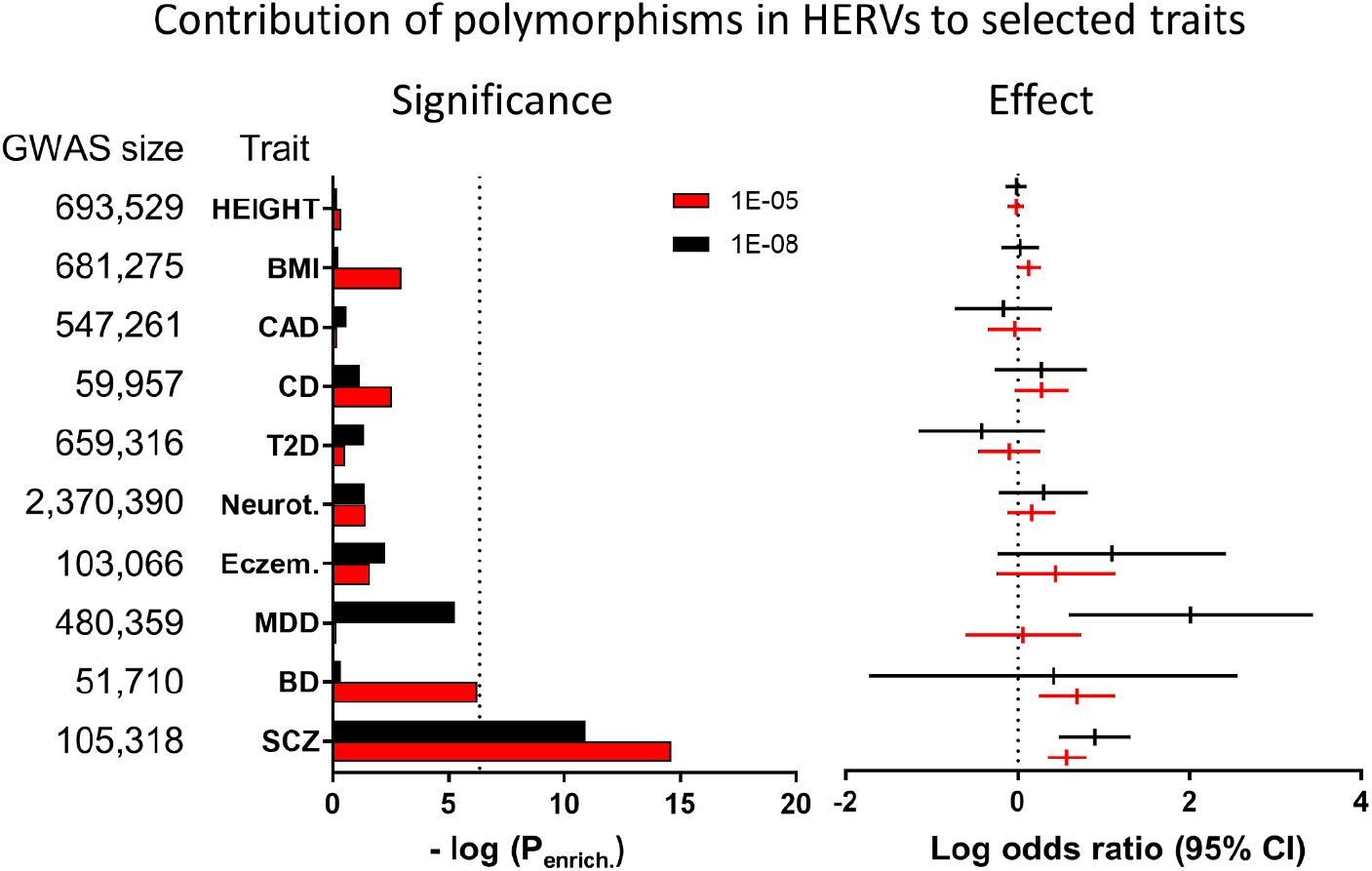
Variants within the retrogenome are enriched with schizophrenia-associated polymorphisms. No association with any other trait was observed (see **Supplemental Table 1** for full results). Calculated using GARFIELD^27^.

### The contribution of locus-specific HERVs and HERV families to schizophrenia susceptibility

We co-localized polymorphisms within individual HERVs and those associated with schizophrenia by GWAS using MAGMA^28^, which calculates gene-level statistics and weighted p-values based on summary statistics whilst adjusting for gene size, single nucleotide polymorphism (SNP) density and linkage disequilibrium. This gene-level enrichment analysis revealed that 148 HERVs from multiple families were significantly enriched for risk variants implicated in schizophrenia, after correcting for the number of HERVs tested (P_corrected_ < 0.05 / 12,389 HERVs in chromosomes 1-22, excluding those at the major histocompatibility locus; **Supplemental Table 2**). The quantile-quantile plot highlights the contribution of many locus-specific HERVs associated with risk, compared to an expected normal distribution (**Figure 2A**), and the Manhattan plot shows their diverse genomic location (**Figure 2B**).

**Figure 2.**
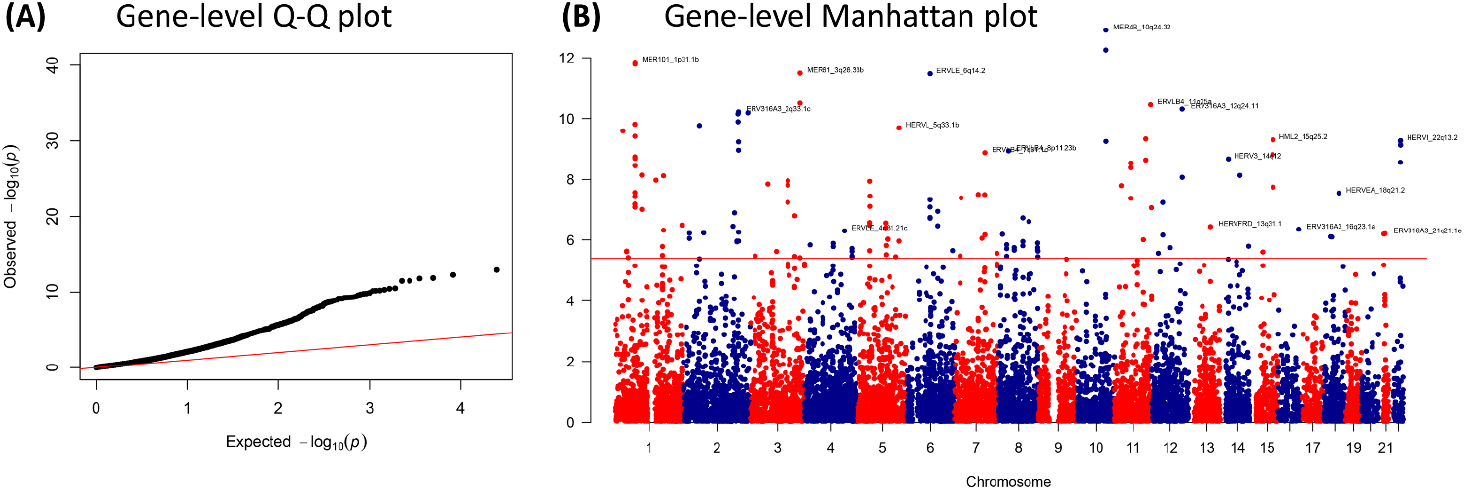
Gene-level enrichment analysis of the schizophrenia GWAS summary statistics using the HERV annotation developed by Bendall and colleagues^20^, calculated using MAGMA^28^. Chromosomes 1-22 only, extended MHC region excluded (chromosome 6, from 26-34 Mb). **(A)** Quantile-quantile plot showing the contribution of several HERVs to schizophrenia genetics compared to an expected normal distribution (red line). **(B)** Manhattan plot showing the location of the HERVs associated with schizophrenia. Plots created using qqman^61^. All enriched HERVs are shown on **Supplemental Table 2**.

To explore the contribution of HERV families towards schizophrenia risk, we investigated whether any of the 60 families defined in the HERV annotation were overrepresented in the list of schizophrenia-associated HERVs using a gene-set enrichment analysis in MAGMA. Each HERV was assigned to a family based on the RepBase model that most closely matched the internal region sequences^20,29^. We observed a nominal association between schizophrenia and the HERVL40 family (P = 0.02, β = 0.14 ± 0.02, SE = 0.07), but this did not survive multiple testing correction (P_corrected_ [for 60 families] > 0.05; **Supplemental Figure 1**). These findings suggest that risk for schizophrenia attributed to the retrogenome may occur via locus-specific sequences, as opposed to entire HERV families.

### HERV expression in the dorsolateral prefrontal cortex

To advance our understanding of HERV expression in the adult brain, we assessed global HERV expression (the retrotranscriptome) in the dorsolateral prefrontal cortex (DLPFC) of 593 post-mortem individuals (N = 593), including 279 unaffected controls, 259 schizophrenia patients, 47 bipolar disorder patients and 8 cases broadly diagnosed with an affective disorder. This was achieved by applying the Telescope pipeline^20^ to the RNA-seq data from the CommonMind Consortium (CMC) dataset. Analysis of these samples revealed that 3,979 HERVs were consistently expressed in the DLPFC according to DESeq2 independent filtering criteria (mean normalized counts = 40.79 [37.8, 43.78]; **Figure 3A**). We performed RT-qPCR to confirm the expression of five arbitrarily selected HERVs in an independent post-mortem cohort of control individuals from the London Neurodegenerative Diseases Brain Bank (N = 10; **Supplemental Material; Supplemental Figure 2**).

**Figure 3.**
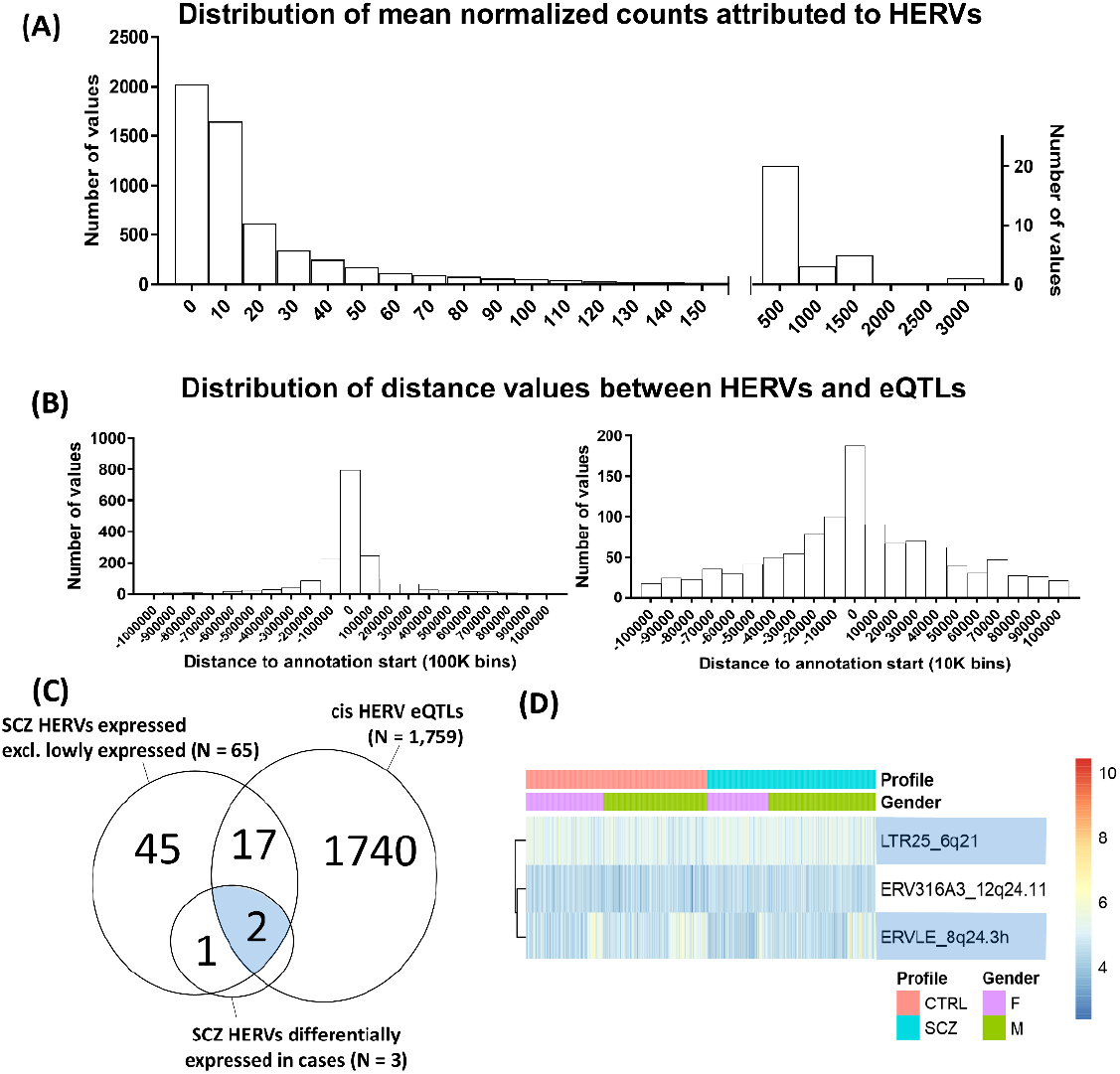
HERV expression in the dorsolateral prefrontal cortex, based on an analysis of the CommonMind Consortium dataset with Telescope. **(A)** Frequency distribution of the mean normalized counts per HERV across samples showing that the majority of HERVs are lowly expressed, according to an analysis of 593 post-mortem brains. Normalized counts do not include adjustments utilized in the analyses. **(B)** Distribution of values representing the distance between the eQTL for a HERV and the start site for that HERV. Left panel shows all data in bins of 100,000, and the right panel only data nearer the start site of the HERVs, in bins of 10,000. A large proportion of eQTLs was located within a 10kb window upstream or downstream the start coordinates of the HERV they regulate. **(C)** Overlap between the 65 schizophrenia-associated HERVs expressed in the brain, the 1,759 HERVs modulated by eQTLs, and the three HERVs enriched for schizophrenia variants and additionally differentially expressed between schizophrenia cases (N = 259) and unaffected individuals (N = 279). **(D)** Heatmap of the six HERVs enriched for schizophrenia variants and further differentially expressed in patients, separated by gender. In blue, HERVs that are modulated by eQTLs, as demonstrated in **Figure 4**.

### Cis-acting HERV eQTLs in the DLPFC

To understand how HERVs are regulated in the brain, we performed an expression quantitative trait loci (eQTL) analysis using all samples (N = 593). Such analysis aims to identify SNPs that explain variation in the expression of HERVs residing in close proximity, to inform about the basic processes responsible for HERV regulation, and to complement our genetic enrichment analyses. The genomic coordinates from the expressed HERVs were remapped to hg19 positions to match genotype information, and only HERVs from chromosomes 1-22 were analyzed (total = 5,349 HERVs, which includes lowly expressed HERVs, according to DESeq2’s internal filtering criteria). This analysis revealed that 1,759 HERVs were regulated in *cis* by 1,622 SNPs located within a 1 Mb window upstream or downstream the start site of the HERV annotation, under the false discovery rate of 5% (q < 0.05). The majority of eQTLs were located within a 10 kb window upstream or downstream of the annotation start site from the HERV they regulate (**Figure 3B**). Of the 148 HERVs that were enriched for schizophrenia variants, we observed that 19 were significantly regulated by eQTLs in the DLPFC (**Supplemental Tables 2 and 3**, highlighted in green; **Figure 3C**).

### Case-control differences in HERV expression

We observed 81 HERVs as differentially expressed between cases (N = 259) and unaffected individuals (N = 279) under the false discovery rate of 5%, independent of a genetic association with schizophrenia (q < 0.05 [corrected for the 3,979 expressed HERVs]; **Supplemental Figure 4**, **Supplemental Table 4**). Our analysis showed that 36 HERVs were downregulated and 45 upregulated in cases, and that expression differences were subtle, with log2 fold-changes of 0.18 ± 0.10 on average.

### Convergent analyses implicate ERVLE_8q24.3h and LTR25_6q21 as robust risk factors for schizophrenia

To make a more informative assessment of HERV expression differences associated with disease status, we first explored whether HERVs identified as genetic risk factors for this disorder were expressed in the DLPFC. Of the 148 HERVs identified via gene-level enrichment analysis, only 65 were consistently expressed across samples according to DESeq2 internal filtering criteria, with mean normalized counts of 57.5, CI 95% [12.59, 102.3]. Of these, we observed that two were significantly upregulated (ERV316A3_12q24.11, log2 fold-change = 0.16, standard error = 0.06; LTR25_6q21, log2 fold-change = 0.09, standard error = 0.04), and one downregulated (ERVLE_8q24.3h; log2 fold-change = −0.09, standard error = 0.04) in schizophrenia cases relative to unaffected controls, under the false discovery rate of 5% (q < 0.05 [corrected for 65 HERVs]; **Figures 3C and D**; **Supplemental Table 5**).

Of the three HERVs enriched for schizophrenia variants and differentially expressed in cases, we observed that two were modulated by eQTLs (**Figures 3C and D**, highlighted in blue), which we initially hypothesized could explain the case-control differences observed. These included **ERVLE_8q24.3h** and **LTR25_6q21**, which were regulated by the top eQTLs, **rs4875048** and **rs174399**, respectively. Importantly, **rs4875048** is associated with schizophrenia and is in linkage disequilibrium with the top association signal at this locus, rs10552126; **rs174399** is in linkage disequilibrium with a risk variant at the locus, rs11153302 (**Table 1**)^14^. Strikingly, no gene within a 1 Mb window upstream or downstream of either of the two top eQTLs was associated with schizophrenia according to a recent analysis by PsychENCODE (http://resource.psychencode.org/)^30^.

**Table 1.** Association of the eQTLs (and variants in linkage disequilibrium) with schizophrenia.

| HERV | SNP | Chromosome | Position | Risk allele | Other allele | Frq cases | OR | SE | P | R<sup>2</sup>‡ | D' ‡ |
| --- | --- | --- | --- | --- | --- | --- | --- | --- | --- | --- | --- |
| ERVLE_8q24.3h | rs4875048 | 8 | 144826671 | A | G | 0.1869 | 1.0553 | 0.011855 | 5.6E-06 | - | - |
|  | rs10552126 | 8 | 144844056 | CAT | C | 0.2018 | 1.05919 | 0.0116 | 7.1E-07 | 0.8954 | 1 |
| LTR25_6q21 | rs174399 | 6 | 111919619 | G* | A | 0.2565 | 1.0025 | 0.010488 | 0.81 | - | - |
|  | rs11153302 | 6 | 111918869 | A | G | 0.506 | 1.0397 | 0.00973 | 6.4E-05 | 0.38 | 1 |

\* This SNP is not associated with schizophrenia, but is in linkage disequilibrium with another variant that is associated with this disorder.
‡ Linkage disequilibrium statistics in relation to the top association signal at the locus which is in LD with the HERV-eQTL.

We observed complex regulatory mechanisms governing expression of these two HERVs. **ERVLE_8q24.3h** was significantly downregulated in cases, but, contrary to what was expected, the risk (A-) allele of **rs4875048** was associated with increased expression of this HERV (β = −0.35, P_β dist._ = 0.002, q = 0.007, **Figure 4A**). Similarly, **LTR25_6q21** was upregulated in patients, but the risk (G-) allele of **rs174399** was associated with reduced expression of this HERV (β = 0.29, P_β dist._ = 0.004, q = 0.02; **Figure 4B**). We investigated the effect of these eQTLs in cases and control individuals separately, which revealed that they influenced HERV expression exclusively in unaffected individuals, suggesting there is a compensatory mechanism in patients counteracting the effects of the eQTLs (**Figures 4A and B**; **Supplemental Table 6**).

**Figure 4.**
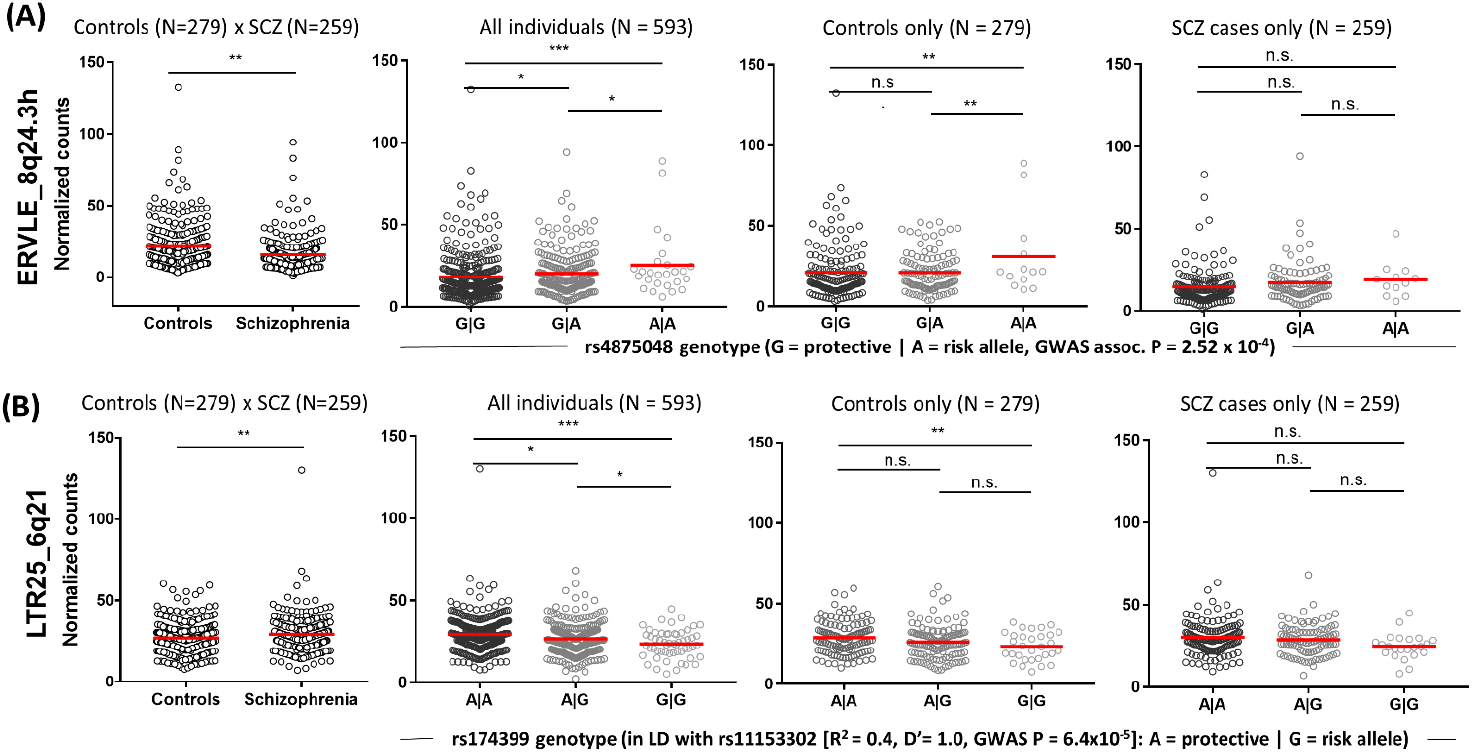
HERVs enriched for schizophrenia variants and differentially expressed between cases and controls, and according to genotype. eQTLs modulated HERV expression exclusively in control individuals. Graphs show normalized counts associated with case-control status (left; Wald tests, **P < 0.01) or per genotype considering all individuals, control individuals or schizophrenia patients only (last three graphs, respectively; ANOVAs followed by pairwise comparisons corrected using the Bonferroni method (*P < 0.05, **P < 0.01, ***P < 0.001; n.s.: not significant; **Supplemental Table 6**). Data shown are for (A) ERVLE_8q24.3h and its top eQTL in the DLPFC, rs4875048, and (B) LTR25_6q21 and its top eQTL, rs174399. Values are uncorrected for the factors and covariates included in the eQTL analysis.

To understand about the regulation of these HERVs during neurodevelopment, we tested their expression in an *in vitro* model of cortical development, which consisted of neural stem cells from the CTX0E16 cell line and cells differentiated for 28 days^31–33^ (**Supplemental Material**, **Supplemental Figure 5**). We observed that both HERVs were differentially regulated during differentiation, suggesting that risk to schizophrenia pertaining these HERVs starts during neurodevelopment. Interestingly, we also observed that both HERVs are located in the antisense strand of *SLC16A10* and *IQANK1* introns, respectively. Moreover, LTR25_6q21 is exclusively present in humans, whereas ERVLE_8q24.3h is shared with primates (**Supplemental Figure 6**), according to the UCSC Genome Browser^34^.

### ERVLE_8q24.3h and LTR25_6q21 are co-regulated with genes implicated in mitochondrial and synaptic function in the adult brain, respectively

To explore the potential function of ERVLE_8q24.3h and LTR25_6q21, we determined the genes that are co-expressed with these HERVs by applying Weighted Correlation Network Analysis (WGCNA)^35^ to the RNA-seq data from schizophrenia patients and controls combined. This systems biology approach is based on the hypothesis that genes within the same co-regulated network share the same function^36^. We observed 19 modules of co-expression in these data (**Supplemental Tables 7 and 8, Supplemental Figure 7**). LTR25_6q21 was assigned to the green module (module membership statistic (MM) = 0.41, P = 3.55 × 10^−23^; gene significance (GS) statistic in relation to case-control status: 0.12, GS P = 0.005), whereas ERVLE_8q24.3h was assigned to the turquoise module (MM = 0.46, P = 4.81 × 10^−29^; GS = −0.15, GS P = 5.14 × 10^−4^). Gene Ontology (GO) analysis of the green module indicates that this gene set is associated with neuronal function, with significant terms including “synapse organization”, “presynapse” and “vesicle-mediated transport in synapse” (q < 0.05, **Figure 5A**, upper panel, **Supplemental Table 9**). The turquoise module, in turn, was significantly associated with mitochondrial function, with terms including the “respiratory chain” and “mitochondrial matrix”, and “NADH dehydrogenase complex assembly” (q < 0.05, **Figure 5A**, lower panel, **Supplemental Table 9**).

**Figure 5.**
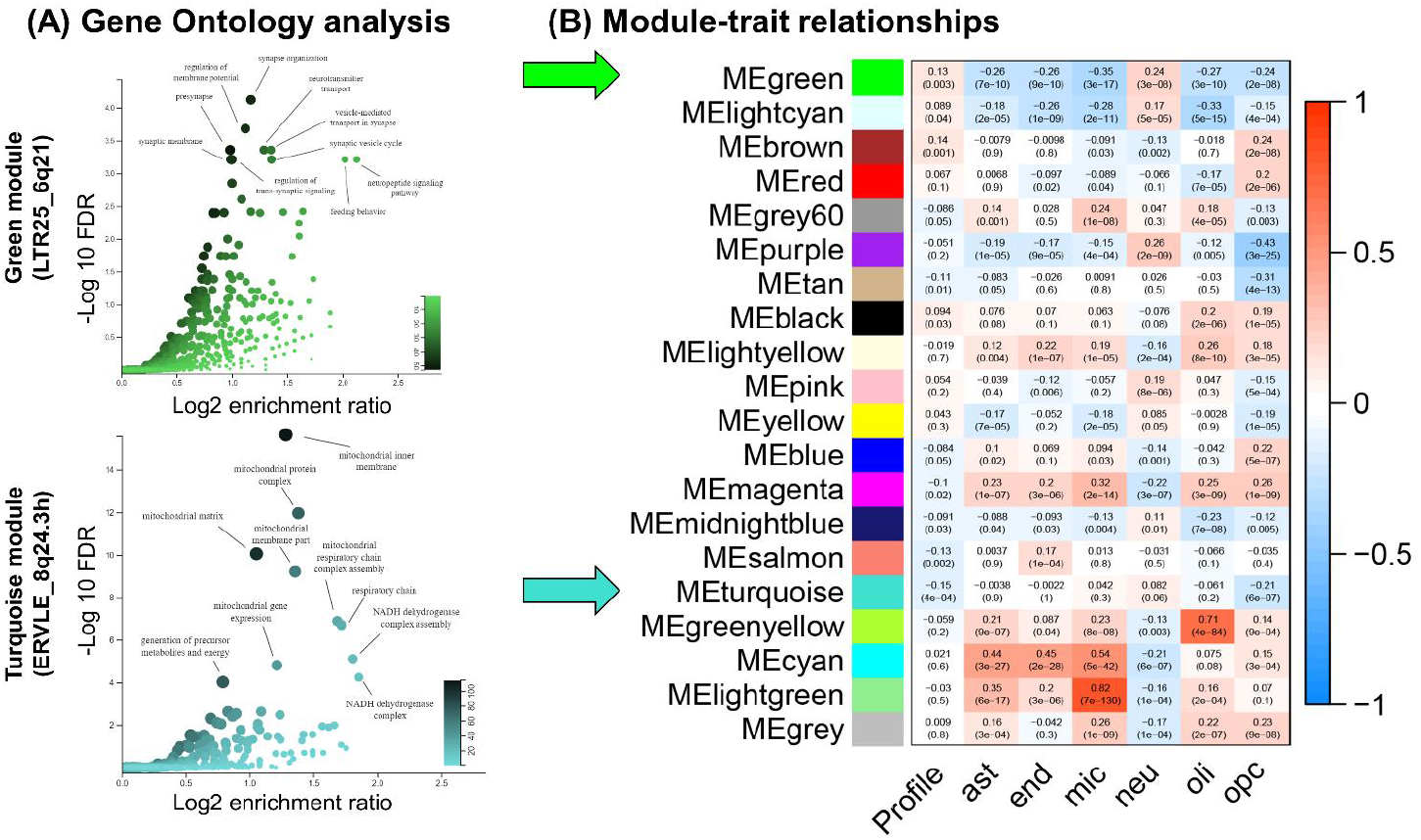
Gene ontology enrichment analysis of the co-expression modules associated with the green (LTR25_6q21) and turquoise modules (ERVLE_8q24.3h), and correlations between modules eigengenes, case-control status (Profile) and coefficients associated with cell counts for major neural cell types. **(A)** LTR25_6q21 belongs to the green module, which is significantly enriched for GO terms associated with neuronal function, as shown in the Volcano plot. ERVLE_8q24.3h, in turn, belongs to the turquoise module, which is significantly enriched for GO terms associated with mitochondrial function. **(B)** The green module is positively associated with neuronal counts, whereas the turquoise module does not correlate strongly with a specific cell type, apart from a negative correlation with oligodendrocyte progenitor cell counts. Each cell of the heatmap contains the Pearson’s r coefficient followed by significance of the correlation (P). Ast: astrocytes, end: endothelial cells, mic: microglia, neu: neurons, oli: oligodendrocytes, opc: oligodendrocyte progenitor cells.

A correlation between the first principal component capturing the variability within each module (module eigengene) and case-control status (‘Profile’) was performed in WGCNA, which revealed that the green module is positively associated with case-control status (r = 0.13, P = 0.003), whereas the turquoise module was negatively associated (r = −0.15, P = 4 × 10^−4^; **Figure 5B**). The coexpression modules were detected assuming a signed network, which means that a positive module-trait correlation entails higher expression of genes in the module in association with disease status, and vice-versa, corroborating the case-control differences observed for each HERV in the previous analysis (**Figures 4A and B**, left panels).

To understand the function of other HERVs associated with schizophrenia from the gene-level enrichment analysis performed using MAGMA (**Supplemental Table 2**), we identified the coexpression modules associated with each enriched HERV that was expressed in the brain (**Supplemental Table 10**). We observed that several of these HERVs were assigned to modules implicated in synaptic organization, including MER41_1p33 and MER41_5p12 alongside LTR25_6q21 in the green module, as well as synaptic function, including HERVL_14q24.2d, HERVL40_8q21.3b, HERVW_6q21c, HML3_5p12a and MER41_2q31.2 in the red module. Interestingly, we observed that these two modules were clustered together in an unsupervised hierarchical clustering analysis performed in WGCNA, corroborating a related or shared function (**Supplemental Figure 8**).

### LTR25_6q21 belongs to a module associated with increased neuronal counts, whereas ERVLE_8q24.3h does not robustly correlate with major neural cell types

We investigated the major cell types associated with each modules eigengene (the first principal component of the module) to infer the cell types associated with ERVLE_8q24 and LTR25_6q21. We estimated the proportion of neural cell types in the RNA-sequencing data by applying the R package BRETIGEA^37^ to the normalized gene counts of the RNA-sequencing data. BRETIGEA estimates the cell type proportions that constitute the sequenced material based on a database of single-cell RNA-sequencing data, ultimately generating coefficients that represent the proportion of astrocytes, microglia, endothelial cells, oligodendrocytes, oligodendrocyte progenitor cells and neurons in the sequenced samples^37^. To infer the cell types associated with each module, we performed correlations between each module and the cell-type proportion coefficients (**Figure 5B**). The module assigned to LTR25_6q21 (green) was significantly correlated with the expression of neuronal markers (r = 0.24, P = 3 × 10^−8^), and negatively associated with the expression of all other cell types (q < 0.05, corrected for the six cell types tested and the two modules of interest), consistent with a role for this module in the regulation of neuronal function, as indicated by the GO analysis. The module assigned to ERVLE_8q24.3h (turquoise) was negatively correlated with the expression of oligodendrocyte progenitor cell markers (r = −0.21, P = 6 × 10^−7^, q < 0.05), but showed no correlation with the other cell types, which may suggest a specific role for this HERV in non-dividing cells. Ultimately, these data indicate that LTR25_6q21 and ERVLE_8q24.3h are part of co-regulated networks implicated in neuronal and mitochondrial function, respectively, suggesting their involvement in these cell functions.

## Discussion

HERVs are ancient viral genetic elements scattered throughout the genome, with previously hypothesized influences on neurodevelopment and risk for schizophrenia. We took a comprehensive approach to reconsider the role of HERVs at the omics level, leveraging on the recent advances in the genomic annotation of HERVs, single-locus resolution quantification, systems biology methods, and population genetic tools.

We investigated the combined contribution of HERVs (the retrogenome) to risk for schizophrenia and other complex polygenic traits, by assessing the overlap between SNPs implicated in these traits and those in genomic locations encompassing HERVs. We were surprised to find that schizophrenia was the only tested trait significantly enriched for common variants within the retrogenome, especially since there is evidence linking HERVs to conditions like amyotrophic lateral sclerosis^7–9^, Crohn’s disease^38^, major depressive disorder and bipolar disorder^10,11^. The fact that the retrogenome is significantly enriched for polymorphisms implicated in schizophrenia suggests that HERVs comprise an important set of risk factors for this disorder, within the ‘non-coding’ genome.

We investigated which HERV families and locus-specific HERVs could be particularly important in moderating risk for schizophrenia at the genetic level by co-localizing risk variants with individual HERV loci from across 60 families. We identified 148 specific HERVs enriched for schizophrenia-associated SNPs. A subsequent gene-set analysis did not find a particular HERV family enriched for schizophrenia variants, further suggesting that disparate HERVs scattered throughout the genome moderate susceptibility rather than whole families. These findings have several implications for the interpretation of previous studies in the literature, and may explain how our results differ from previous data implicating other HERVs or HERV families as risk factors, which may not have been identified here. Most HERV expression research to-date has been performed using microarrays, antibodies or RT-qPCR probes, which do not provide sufficient specificity to assess single HERVs^7–11^. Therefore, previous studies may have captured the expression of multiple HERVs concomitantly due to their repetitive sequences. Based on our findings, it is unlikely that individual HERVs (even from within the same family) contribute equally to risk. The HERV annotation developed by Bendall and colleagues^20^, as well as the Telescope pipeline, and the application of modern population genetic methods, enabled us to revisit the role HERVs play in relation to schizophrenia risk on a genome-wide scale, and allow for more robust inferences regarding their etiological relevance.

Next, to better understand HERV expression regulation in the brain, we analyzed RNA-sequencing data from the CommonMind Consortium, which confirmed widespread HERV expression in the adult brain and cis-regulatory mechanisms. We found 3,979 HERVs actively expressed in a key brain area linked to schizophrenia pathophysiology, the DLPFC^39^. We further identified 1,759 HERVs that were regulated by 1,622 short-range (cis-) eQTLs in the DLPFC. The identification of eQTLs that impact HERV expression informs us about the basic processes responsible for HERV regulation, and can be useful for the interpretation of GWAS findings in the context of the retrogenome^40,41^. In addition, these results suggest that SNPs within HERVs are not simply affecting the expression of neighboring protein-coding genes via their LTRs, rather it demonstrates that common genetic variation impacts locally on HERV expression.

We used a complementary set of analyses to identify the most robust HERVs implicated in schizophrenia. Two HERVs, **ERVLE_8q24.3h** and **LTR25_6q21**, identified from the gene-level enrichment analysis, were found to be regulated by schizophrenia-associated eQTLs, and were found to be differentially expressed in the DLPFC of patients and in an *in vitro* model of cortical neurodevelopment. These findings suggest a potential risk mechanism for schizophrenia that starts during neurodevelopment and persists through to adulthood, as observed for schizophrenia risk genes such as *NT5C2*, *AS3MT*, and *BORCS7*^33,42^. Importantly, we observed a complex regulation of both HERVs, whereby the lead eQTLs only exerted their regulatory effects in unaffected individuals. This suggests that compensatory mechanisms (e.g. epigenetic alterations) may be acting to correct for the effects of the risk variants on the expression of these HERVs in the DLPFC of schizophrenia patients.

We also describe here, for the first time, the co-regulation of several HERVs with known genes in the adult brain, and the GO terms associated with each co-expression module. Our findings suggest that LTR25_6q21 is implicated in neuronal function, whereas ERVLE_8q24.3h is involved in mitochondrial function. WGCNA has been successfully used to predict the biological function of unknown genes or non-coding RNAs in different organisms^43,44^, and to identify clinically relevant cell types when in combination with cell-type deconvolution analysis^45^, and thus represents a powerful approach to functionally characterize the HERVs expressed in the brain. For a long time HERVs were assumed to be mere regulatory DNA sequences, but the discovery of their expression and co-regulation with several other genes implicated in multiple biological processes in the brain, ranging from neuronal, glial and mitochondrial regulation to splicing and cell motility (**Supplemental Table 9**), is a landmark for HERV research, and adds an extra layer of complexity to our understanding of human neurobiology.

There are limitations to this study which should be acknowledged. Schizophrenia is a highly polygenic, heterogeneous disorder, and as such large sample sizes are required for appropriate comparisons. The CommonMind Consortium provides the largest and best characterized cohort of schizophrenia cases and unaffected individuals with RNA-sequencing data to-date, but it might be underpowered for case-control comparisons considering the heterogeneity of schizophrenia. Nevertheless, we complemented case-control comparisons with genomic and eQTL analyses to provide additional insights. Another limitation to our study is that it investigated RNA-sequencing data from bulk DLPFC tissue only, which is composed of a heterogeneous mixture of several types of neurons and glial cells, and it is possible that HERVs expressed in particular cell types, or in other brain regions, are more relevant to risk^46^. To address this, we performed cell type deconvolution using BRETIGEA to determine cell-type specific effects, but ultimately the analysis of data from other brain areas, developmental time points and single-cell datasets has the potential to reveal important insights about the etiology of schizophrenia in relation to HERV expression. In addition, our post-mortem and *in vitro* work suggest HERV expression is important in the DLPFC and during its development, but we still do not know the function of these HERVs. To infer function we performed WGCNA which provides insight into which processes risk HERVs moderate, but future functional studies are required to definitively characterize how HERVs, particularly ERVLE_8q24.3h and LTR25_6q21, influence the transcriptome, neural stem cell proliferation or neuronal differentiation in the context of schizophrenia risk, as is currently being investigated in relation to protein-coding risk genes^32,33,47,48^.

The development of a retrogenome annotation, and advances in modern population genetic methods and transcriptomic tools, now allows us to investigate HERVs at the omics level, in the context of risk for many biological traits. Our work studying the role of HERVs in the brain, and their relationship to schizophrenia ignites a new, provocative line of thought implicating HERVs as biological risk factors for schizophrenia and confirms that these previously assumed ‘dormant’ sequences in the brain may not be dormant after all.

## Online Methods

We used a combination of gene expression, genetic and *in vitro* analyses to identify the most robust HERVs implicated in schizophrenia risk (**Figure 6**). Further details are provided in the **Supplemental Material**.

**Figure 6.**
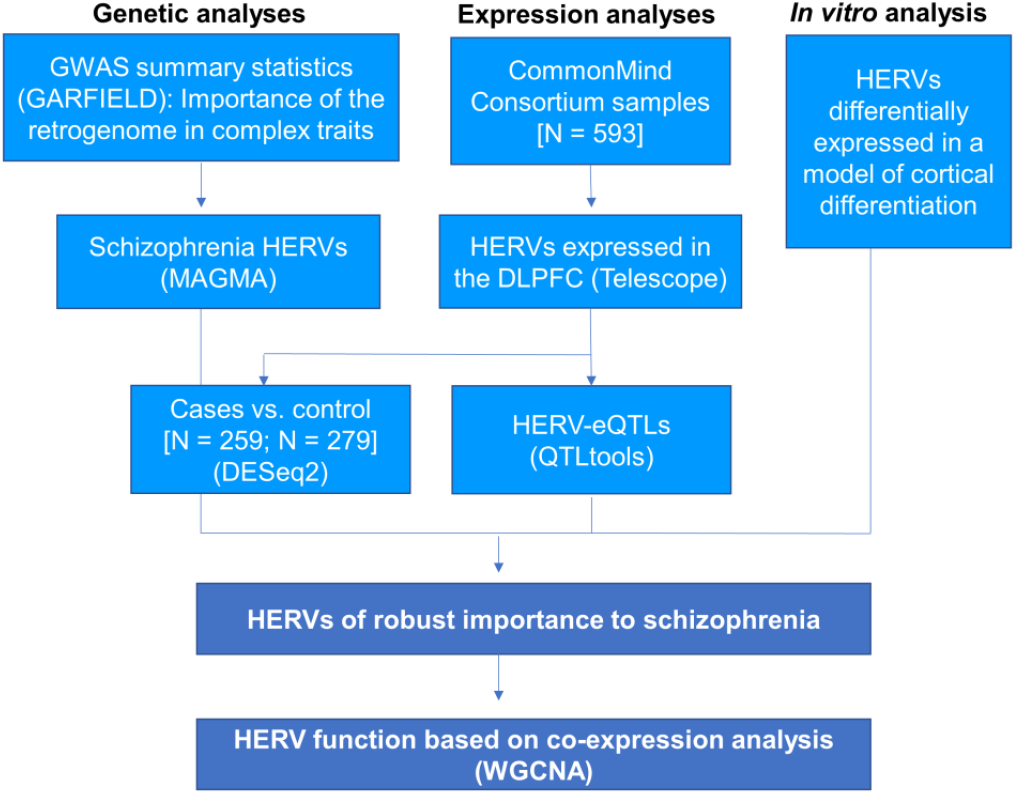
Analysis strategy. We performed a series of analyses using post-mortem brain RNA- sequencing data, genetic enrichment analyses and *in vitro* cortical differentiation data to identify HERVs of robust importance in schizophrenia. Bioinformatic tools used are indicated in parentheses. DLPFC: dorsolateral prefrontal cortex.

### Genetic enrichment analyses

We estimated the contribution of genetic polymorphisms within the retrogenome towards risk of developing multiple traits using GARFIELD^27^. We downloaded summary statistics from well-powered genome-wide association studies, including of schizophrenia (N = 105,318 individuals)^14^, height (N = 693,529)^49^, body mass index (N = 681,275)^49^, coronary artery disease (N = 547,261)^50^, Crohn’s Disease (N = 59,957)^51^, type 2 diabetes (N = 659,316)^52^, neuroticism (N = 2,370,390)^53^, eczema (N = 103,066)^54^, major depressive disorder (N = 480,359)^55^, bipolar disorder (N = 51,710)^56^, Alzheimer’s disease (N = 74,046)^57^, attention deficit hyperactivity disorder (N = 53,293)^58^, amyotrophic lateral sclerosis (N = 36,052)^59^, and autism spectrum disorder (N = 46,350)^60^, which analyzed European cohorts only. GARFIELD performs greedy pruning of SNPs in GWAS summary statistics (those in linkage disequilibrium, with R^2^ > 0.1), and quantifies enrichments using odds ratios, assessing their significance by employing generalized linear model testing, controlling for minor allele frequency, and number of linkage disequilibrium proxies (R^2^ > 0.8). Linkage disequilibrium and allele frequency information were calculated based on the UK10K study. Enrichments were calculated based on summary statistics from each trait using two P_association_ thresholds: P < 5 × 10^−8^, to test the enrichment of HERVs within genome-wide significant variants; and a more relaxed threshold, P < 5 × 10^−5^, to allow signal capture in less powered GWAS. The enrichment significance was corrected for the number of tests performed [14 traits and two P-value thresholds tested]. For consistency with the GWAS data, the HERV annotation used in the expression analysis (hg38) was remapped to hg19 coordinates using liftOver^34^.

To identify locus-specific HERVs and potential HERV families associated with schizophrenia, we used MAGMA 1.07b^28^. Briefly, MAGMA calculates gene-level enrichment by generating a gene-wide statistic from summary statistics, adjusting for gene size, variant density, and linkage disequilibrium using the 1000 Genomes Phase 3 European reference panel. SNPs from the summary statistics were assigned to HERVs using an annotation window of 10 kb upstream and downstream of each HERV (as suggested by the authors)^28^. A Bonferroni correction was applied to identify significantly enriched HERVs (P_cut-off_ < 4.03 × 10^−6^ [0.05 / 12,393 HERVs in chromosomes 1-22, excluding the major histocompatibility locus]). Q-Q and Manhattan plots were generated using qqman 0.1.4^61^. Gene-set enrichment analysis was additionally performed using MAGMA, to test whether HERVs associated with schizophrenia in the previous step were enriched for any of the 60 HERV families (excluding HERVs located in sex chromosomes or at the MHC locus – chromosome 6: 26 – 34 Mb).

### The CommonMind Consortium dataset

To identify HERV expression differences in schizophrenia patients, or HERV expression quantitative trait loci (eQTL) in the dorsolateral prefrontal cortex, we analyzed RNA-sequencing data from the CommonMind Consortium (release 1.0, N = 593 individuals, https://doi.org/10.7303/syn2759792)^39^. This dataset consisted of dorsolateral prefrontal cortex (DLPFC) samples from 279 unaffected individuals, 259 schizophrenia cases, 47 bipolar disorder patients and 8 cases broadly diagnosed with an affective disorder. Access to this dataset, which includes expression, genotype and clinical data, was granted under a Material Transfer Agreement with the NIMH Repository and Genomics Resources. Briefly, autopsy samples from the Mount Sinai NIH Brain Bank and Tissue Repository, the University of Pennsylvania Brain Bank of Psychiatric illnesses and Alzheimer’s Disease Core Center, and The University of Pittsburgh Brain Tissue Donation Program, were sent to the Icahn School of Medicine at Mount Sinai for nucleic acid isolation and sequencing. Individuals were diagnosed according to the Diagnostic and Statistical Manual of Mental Disorders, 4th Edition, as determined in consensus conferences after review of medical records and interviews of family members and care providers. Total RNA was extracted from autopsy tissue using the RNeasy kit (QIAGEN, Hilden, Germany). Ribosomal RNA was depleted using the Ribo-Zero Magnetic Gold kit (Illumina, San Diego, California, United States), libraries were constructed using the TruSeq RNA Sample Preparation Kit v2 (Illumina), and samples were sequenced on an Illumina HiSeq 2500. For whole-genome genotyping, DNA was extracted using the DNeasy Blood and Tissue Kit (QIAGEN) according to the manufacturer’s protocol, and samples were genotyped using Illumina Infinium HumanOmniExpressExome 8 1.1b chips. Further details on quality control and sample processing are described in Fromer and colleagues^39^.

### RNA-sequencing data processing and HERV expression quantification

Bam files containing mapped and unmapped RNA-sequencing reads aligned to the human reference genome (hg19) using TopHat 2.0.9 and Bowtie 2.1.0, were downloaded to the King’s College London High Performance Computer Cluster Rosalind, using the synapse client (1.7.5). Bam files were merged, and fastq files were extracted using samtools 1.5^62^ and the flag ‘−F 0×100’. Trimmomatic 0.38^63^ was used to prune Illumina adaptors, low quality bases (leading/trailing sequences with phred score < 3, or those with average score < 15 every four bases), or reads below 36 bases in length. Trimmed reads were mapped to the human genome hg38 using bowtie2^64^ and the parameters --very-sensitive-local, −k 100, and --score-min L,0,1.6. Subsequently, Telescope 1.0.2 was used to quantify expression of 14,968 HERVs (annotation version hg38), which we defined as the retrogenome^20^. We analyzed HERVs with counts > 10 across 4 samples at least, to avoid inflation driven by lowly expressed elements. Case-control expression differences (N = 538 individuals in total) were calculated in R^65^ using Wald tests in DESeq2^66^, where data was normalized (median of ratios) and controlled for the main confounders of gene expression estimated by Fromer and colleagues^39^, which included institution of sample origin, RNA integrity number, gender, post-mortem interval, age (determined in five bins: #1 = 13-29 years, #2 = 30-49 years, #3 = 50-69 years, #4 = 70-89 years, #5 = 90+ years), as well as the first five population covariates estimated using multidimensional scaling in PLINK 1.9^67^, and the first ten hidden HERV expression confounders estimated using sva^68^, considering schizophrenia and unaffected individuals only.

### Whole-genome genotype data processing

Markers with zero alternate alleles, genotyping call rate < 0.98, Hardy-Weinberg P < 5 × 10^−5^, or individuals with genotyping call rate < 0.90, were removed from the analysis, as described by Fromer and colleagues^39^. PLINK files were generated containing genotype information for 958,178 variants for the 593 subjects. Marker alleles were phased to the forward strand, and ambiguously stranded markers were removed. Additional genotype information was imputed from the 1000 Genomes Phase 1 reference panel using minimac3 and Eagle v2.3 phasing with the Michigan Imputation Server (https://imputationserver.sph.umich.edu/index.html). Genotype information from the 22 autosomes was concatenated using bcftools 1.9 (https://samtools.github.io/bcftools/bcftools.html), non-single nucleotide polymorphisms (SNPs) were excluded, as well as sites with an imputation R^2^ < 0.8, minor allele frequency < 0.05, or Hardy-Weinberg P < 5 × 10^−5^.

### eQTL analysis

Normalized HERV counts per sample were obtained using DESeq2 (N = 593), and HERV expression was tested for the effect of genotype at all variants located within a 1 Mb window upstream or downstream from the annotation start site of each HERV using QTLtools^69^, according to the authors’ manual. We covaried for the effect of case-control status, institution of sample origin, RNA integrity number, gender, post-mortem interval, age (five bins, as described previously), the first five population covariates, and ten hidden expression confounders estimated using sva^68^ (**Supplemental Figure 3**). eQTL P-values were corrected through estimation of a beta distribution using a minimum of 1,000 permutations and maximum of 10,000, and were further corrected for the number of HERVs tested using the false discovery rate method (q < 0.05).

### Weighted Correlation Network Analysis (WGCNA)

WGCNA is a systems biology approach that enables the identification of co-expressed genes in transcriptomic data, which we used here to identify the genes co-expressed with schizophrenia HERVs in order to infer their biological function^35^. We used this tool to construct a signed network consisting of HERVs and genes, which was created based on an adjacency matrix that informs about the co-expression similarity observed between all pairs of genes and HERVs in the expression data (i.e. genes and genes, genes and HERVs, HERVs and HERVs). Normalized HERV and gene counts were variance-stabilized in DESeq2, and were further adjusted for all confounders previously described, using the *removebatcheffect* function in limma^70^. To achieve this, we combined gene and HERV counts obtained from the brain of 538 individuals (279 unaffected individuals and 259 schizophrenia cases), and filtered out genes and HERVs that were lowly expressed, i.e. those with < 10 counts in < 80% of samples), as these can drive spurious correlations^35^. The normalized counts were variance stabilized transformed using DESeq2 and adjusted for institution of sample origin, gender, case-control status, age bins, post-mortem interval, the first five population dimensions (estimated in plink), and RIN, using limma^70^. WGCNA identifies modules by applying hierarchical clustering to the adjacency matrix, further filtering spurious relationships through the application of a topological overlap approach. We used an R^2^ cut-off of 0.8, which corresponds to a β = 12, to construct the network. Each module was assigned a color, and genes or HERVs not belonging to any module were assigned to the gray module. The relationship between modules and specific cell types was tested based on the correlation between the module eigengenes (ME), defined as the first principal component of the module, and cell count estimates, as described below. We applied the false discovery rate method to correct for the module-cell type associations (q < 0.05). Plots were generated by WGCNA.

### Cell type estimates and module correlations

We performed a BRain cEll Type specIfic Gene Expression Analysis (BRETIGEA)^37^ to estimate the abundance of major neural cell types in the 538 samples analyzed by WGCNA. Briefly, BRETIGEA uses expression data from single cell RNA-sequencing data sets to identify the proportion of astrocytes, microglia, endothelial cells, oligodendrocytes, oligodendrocyte progenitor cells and neurons, in bulk brain gene expression data. More specifically, this tool uses a panel of 50 well-established cell type-specific markers to generate coefficients that represent the proportion of each cell type per sample, which were tested for association with each module.

### Gene Ontology (GO) analyses

We performed GO analyses using the WEB-based GEne SeT AnaLysis Toolkit (Webgestalt)^71^ to identify the function of the genes co-regulated with the schizophrenia HERVs in the brain, and thus infer the potential function of these HERVs. All genes inputted to WGCNA were used as background (reference) gene set. We used the false discovery rate method to correct for the GO enrichment analyses within Webgestalt (q < 0.05) and report up to 10 significant GO terms per module. Volcano plots were generated in Webgestalt.

### Statistical analysis and data visualization

The co-localization of GWAS-supported variants with the retrogenome was calculated using linear regressions in GARFIELD^27^, and the gene-level and gene-set enrichment analyses were calculated in MAGMA^28^. Findings were corrected for multiple testing using the Bonferroni method. The case-control HERV expression differences (N = 538 individuals) and effects of eQTLs on HERV expression (N = 593 individuals) were calculated, respectively, using Wald tests in DESeq2, and stepwise linear regressions in QTLtools, respectively. These were corrected using the false discovery rate (q < 0.05), a more permissive multiple testing correction method, to increase our detection power. The effect of genotype on specific HERVs within cases and control groups, separately or combined, was calculated using linear regressions in IBM Statistics SPSS 25 (IBM Corp., Armonk, NY, United States). Other analyses were performed in R^65^. Graphs were generated in R or Graph Pad Prism 7 (GraphPad Software, San Diego, CA, United States).

## Acknowledgements

The work was supported in part by US National Institutes of Health grants: CA206488 (DFN), AI076059 (DFN) and UL1TR001876 (KAC). This work was also supported by a grant from the Coordenação de Aperfeiçoamento de Pessoal de Nível Superior (CAPES, Brazil, Science without Borders award no. BEX1279/13-0) and an NIHR Maudsley Biomedical Research Centre Career Development Award to RRRD. TRP is funded by a Medical Research Council (MRC) Skills Development Fellowship (MR/N014863/1). This study represents independent research part funded by the NIHR-Wellcome Trust King’s Clinical Research Facility and the National Institute for Health Research (NIHR) Biomedical Research Centre at South London and Maudsley NHS Foundation Trust and King’s College London. The views expressed are those of the authors and not necessarily those of the NHS, the NIHR or the Department of Health and Social Care. Data for this publication were obtained from NIMH Repository & Genomics Resource, a centralized national biorepository for genetic studies of psychiatric disorders. Data were generated as part of the CommonMind Consortium supported by funding from Takeda Pharmaceuticals Company Limited, F. Hoffman-La Roche Ltd and NIH grants R01MH085542, R01MH093725, P50MH066392, P50MH080405, R01MH097276, RO1-MH-075916, P50M096891, P50MH084053S1, R37MH057881, AG02219, AG05138, MH06692, R01MH110921, R01MH109677, R01MH109897, U01MH103392, and contract HHSN271201300031C through IRP NIMH. Brain tissue for the study was obtained from the Mount Sinai NIH Brain and Tissue Repository, the University of Pennsylvania Alzheimer’s Disease Core Center, the University of Pittsburgh NeuroBioBank and Brain and Tissue Repositories, and the NIMH Human Brain Collection Core. CMC Leadership: Panos Roussos, Joseph Buxbaum, Andrew Chess, Schahram Akbarian, Vahram Haroutunian (Icahn School of Medicine at Mount Sinai), Bernie Devlin, David Lewis (University of Pittsburgh), Raquel Gur, Chang-Gyu Hahn (University of Pennsylvania), Enrico Domenici (University of Trento), Mette A. Peters, Solveig Sieberts (Sage Bionetworks), Thomas Lehner, Stefano Marenco, Barbara K. Lipska (NIMH). Tissue samples used in RT-qPCR experiments were supplied by The London Neurodegenerative Diseases Brain Bank, which receives funding from the Medical Research Council and as part of the Brains for Dementia Research programme, jointly funded by Alzheimer’s Research UK and Alzheimer’s Society. We thank Professor Cathryn Lewis for her comments on the manuscript and Daniel Bean (King’s College London) for his assistance with coding. We also thank Richard “Brad” Jones, Mario Ostrowski, and Nathaniel Bachtel for their comments on this manuscript.

## Author contributions

Study design: TRP, RRRD. Performed analyses and experiments: RRRD, TRP. Contributed reagents, biological material, revised the manuscript: MLB, MM, CEO, GAB, SS, CT, GRT, KAC, DPS, DFN. Wrote the paper: RRRD, TRP.

## Data availability

Telescope and the HERV annotation are available at http://github.com/mlbendall/telescope. Access to the CommonMind Consortium dataset can be requested to the NIMH Repository & Genomics Resource via https://www.nimhgenetics.org/resources/commonmind.

