## Supplemental Information for "Schizophrenia risk from locus-specific human endogenous retroviruses"

#### *Supplemental Figures*

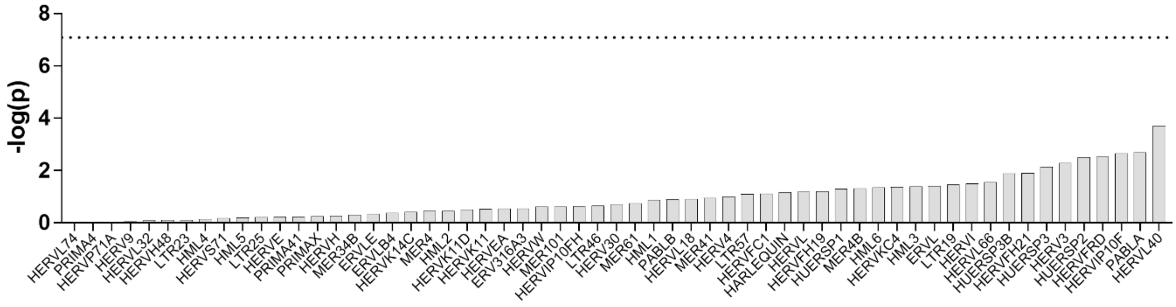


**Supplemental Figure 1.** Individual HERV families are not associated with schizophrenia genetics. Enrichments were calculated in MAGMA. Dotted line represents the natural logarithm of the P cut-off (P = 0.05, corrected for 60 tests = $8.3 \times 10^{-4}$).

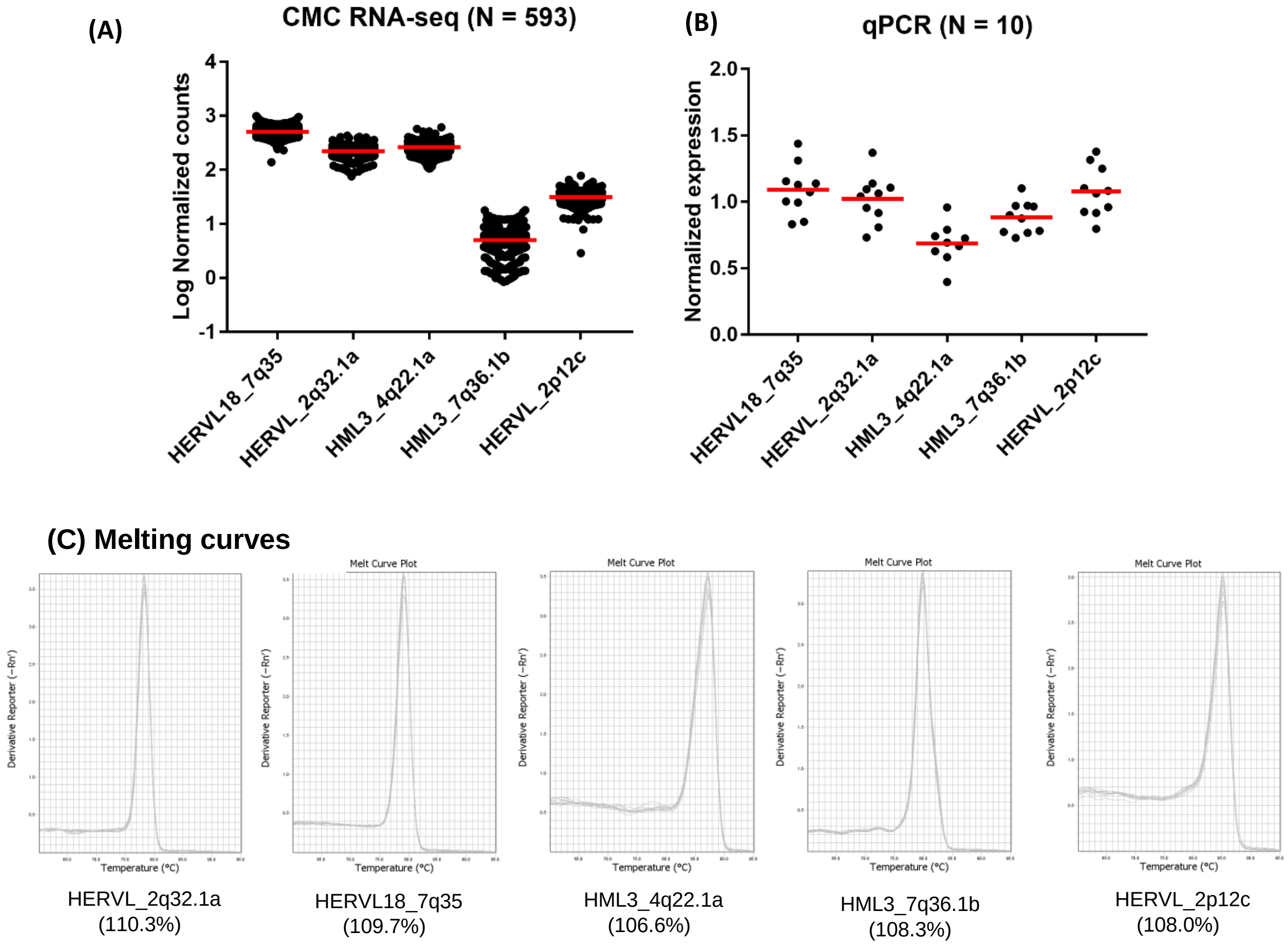


**Supplemental Figure 2.** HERVs identified in the RNA-sequencing data from the CommonMind Consortium (human DLPFC) were confirmed to be expressed at equivalent levels in an independent cohort of post-mortem individuals, using RT-qPCR. **(A)** Expression (log-transformed, normalized counts) of five arbitrarily selected HERVs detected as expressed in the DLPFC, based on analysis of the CMC dataset. **(B)** RT-qPCR primers were designed to amplify these HERVs, to confirm their expression in the DLPFC of an independent group of control individuals from the London Neurodegenerative Diseases Brain Bank (N = 10; see **Supplemental Materials** below). Expression was normalized to a standard curve of five points, and is relative to the geometric mean of the normalized expression of three housekeeping genes (*ACTB*, *SDHA*, and *ALG2*). **(C)** Melting curves with the primers utilized corroborate their specificity in PCR. Values in brackets represent primer efficiency, calculated based on a standard curve of five 1:2 dilution points.

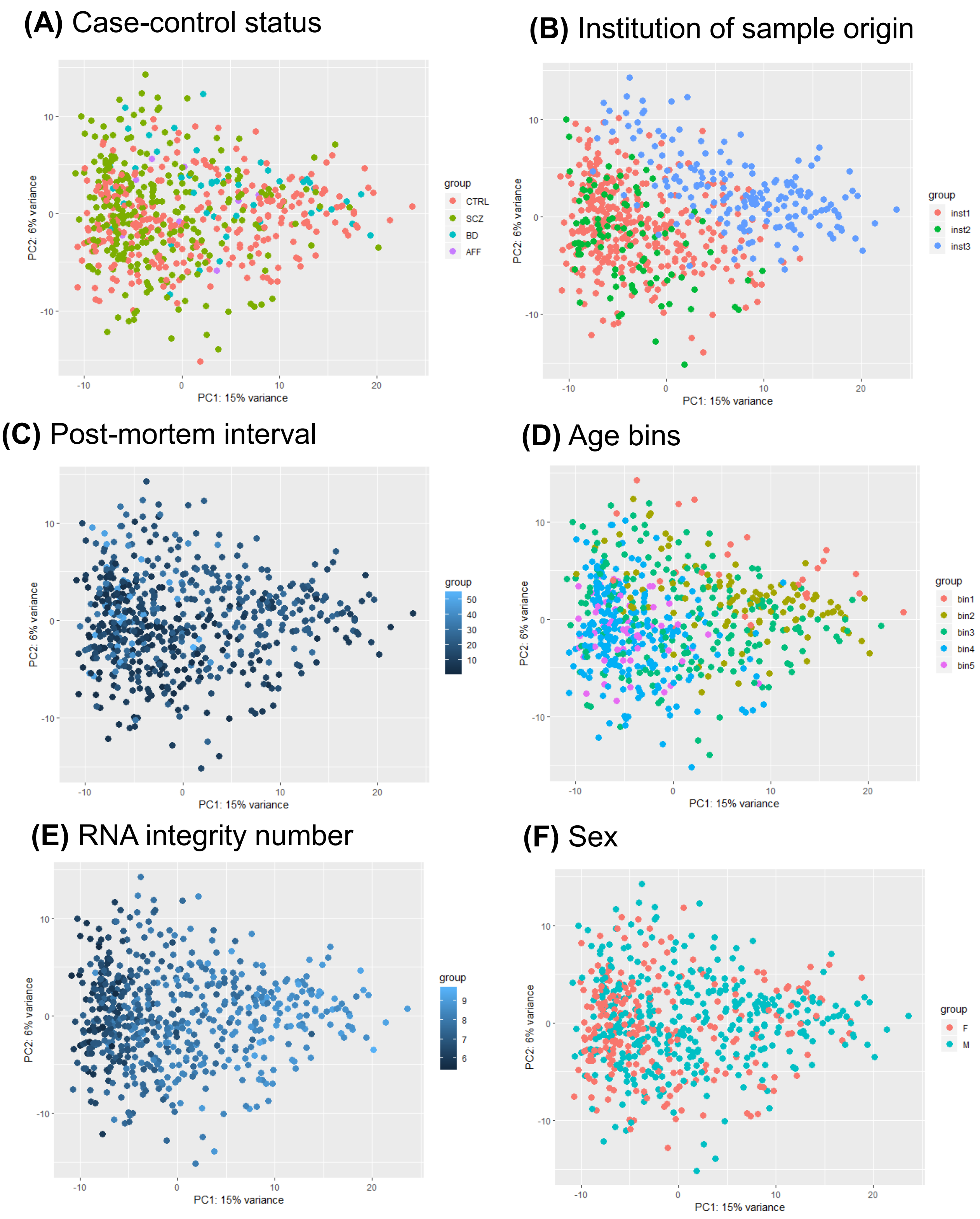


**Supplemental Figure 3.** Principal component plots of HERV expression colored according to the factors observed by Fromer and colleagues [1] to impact on gene expression in the CommonMind Consortium dataset: **(A)** case-control status, **(B)** institution of sample origin, **(C)** post-mortem interval, **(D)** age, **(E)** RNA integrity number, and **(F)** sex. PCAs were calculated using the full cohort (N = 593), and expression data from chromosomes 1-22 only.

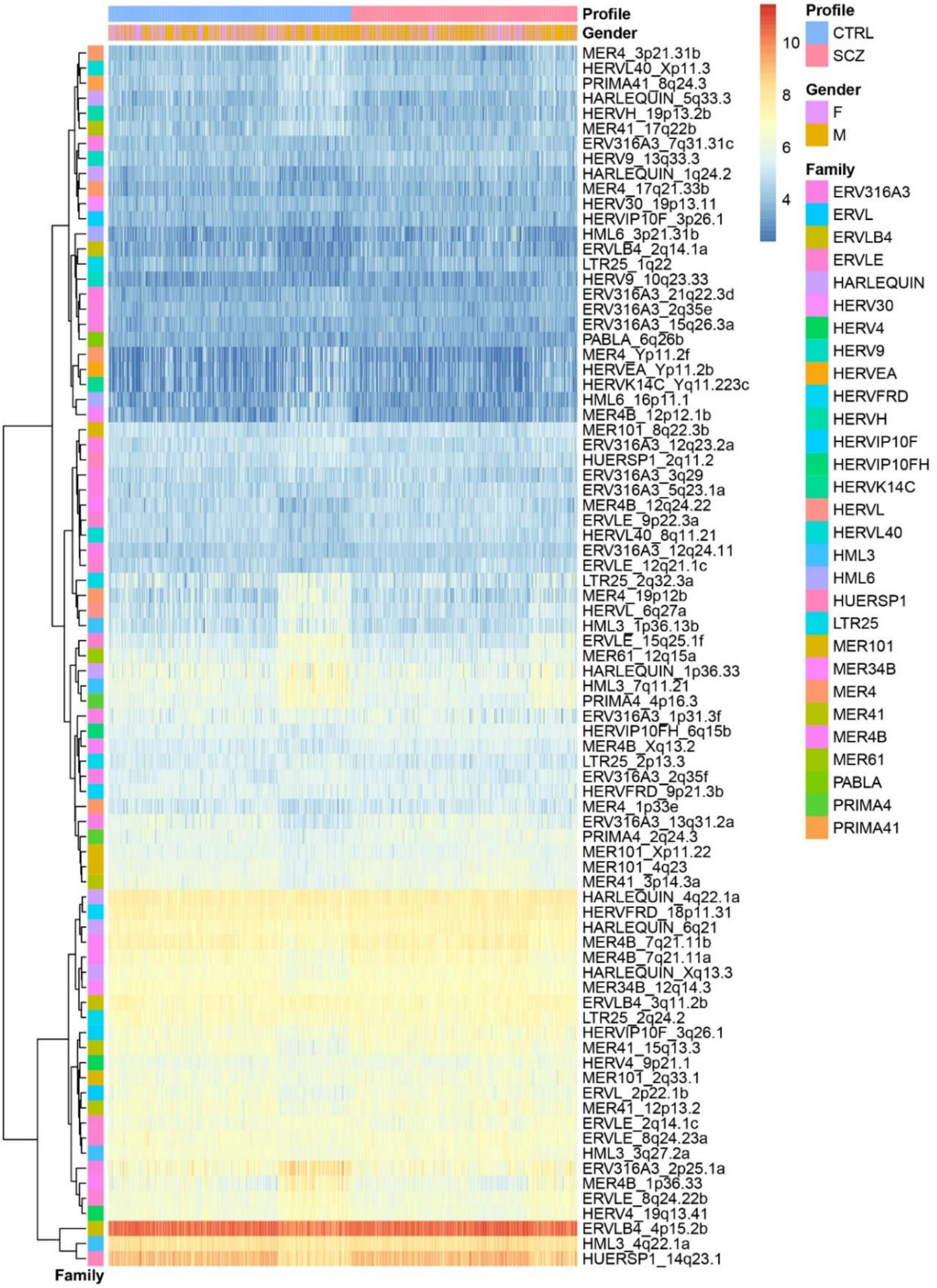


**Supplemental Figure 4.** HERV expression differences observed in the DLPFC of post-mortem schizophrenia patients and controls individuals, under the false discovery rate of 5% (Wald tests, $q < 0.05$).

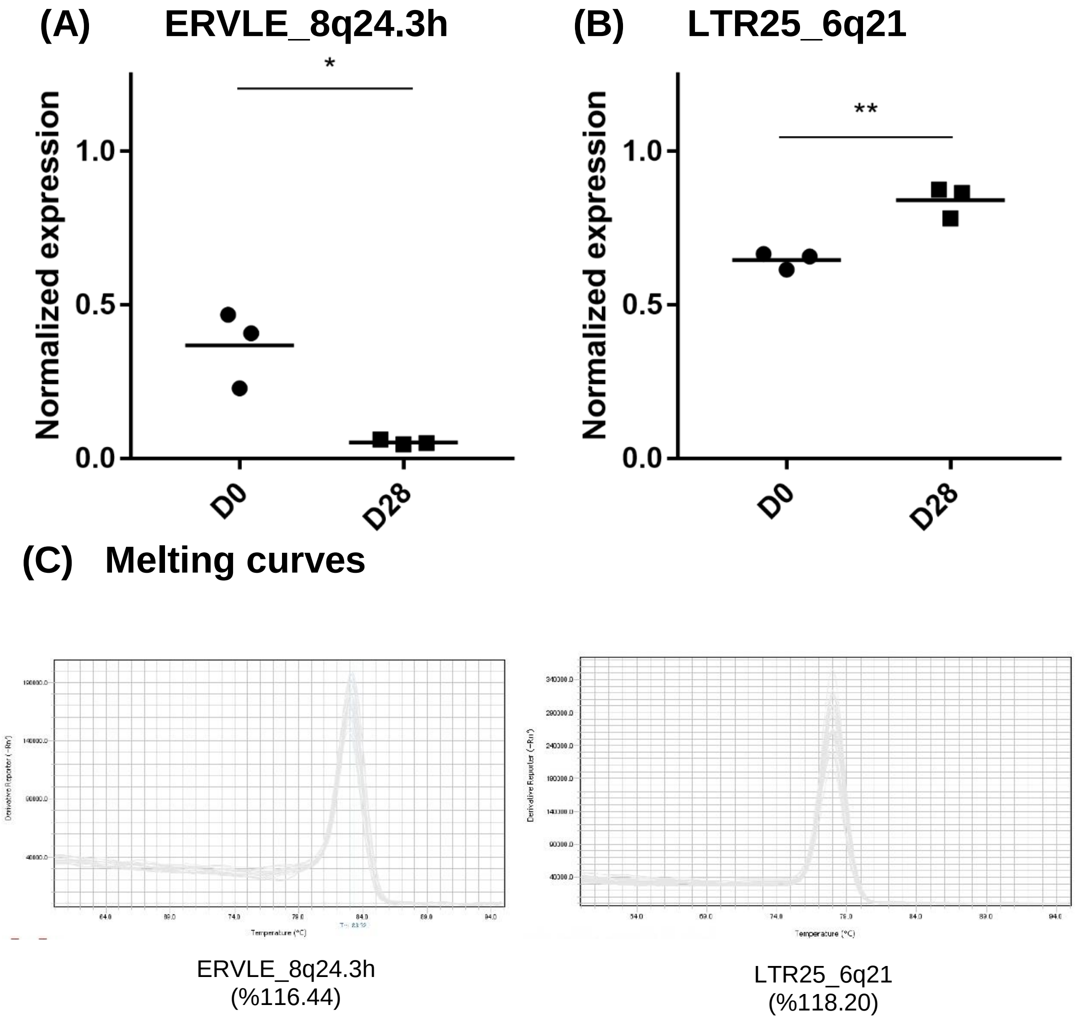


**Supplemental Figure 5.** The HERVs implicated in schizophrenia are neurodevelopmentally regulated, as indicated by analysis in an *in vitro* model of cortical development [2-4]. HERV expression was compared in neural stem cells (D0) from the CTX0E16 cell line, and cells differentiated for 28 days (D28) (Supplemental Materials). **(A)** ERVLE_8q24.3h was substantially downregulated after differentiation (log 2 fold-change = -2.80; t-test, $t(4) = 4.38$; $P = 0.012$; P corrected (for 2 tests) = 0.024; N = 3 biological replicates per condition), whereas **(B)** LTR25_6q21 was upregulated (log 2 fold-change = 0.37; t-test, $t(4) = 5.81$; $P = 0.004$; P corrected (for 2 tests) = 0.008; N = 3). * $P < 0.05$, ** $P < 0.01$. **(C)** Melting curves with the primers utilized corroborate their specificity in PCR. Values in brackets represent primer efficiency, calculated based on a standard curve of five 1:2 dilution points.

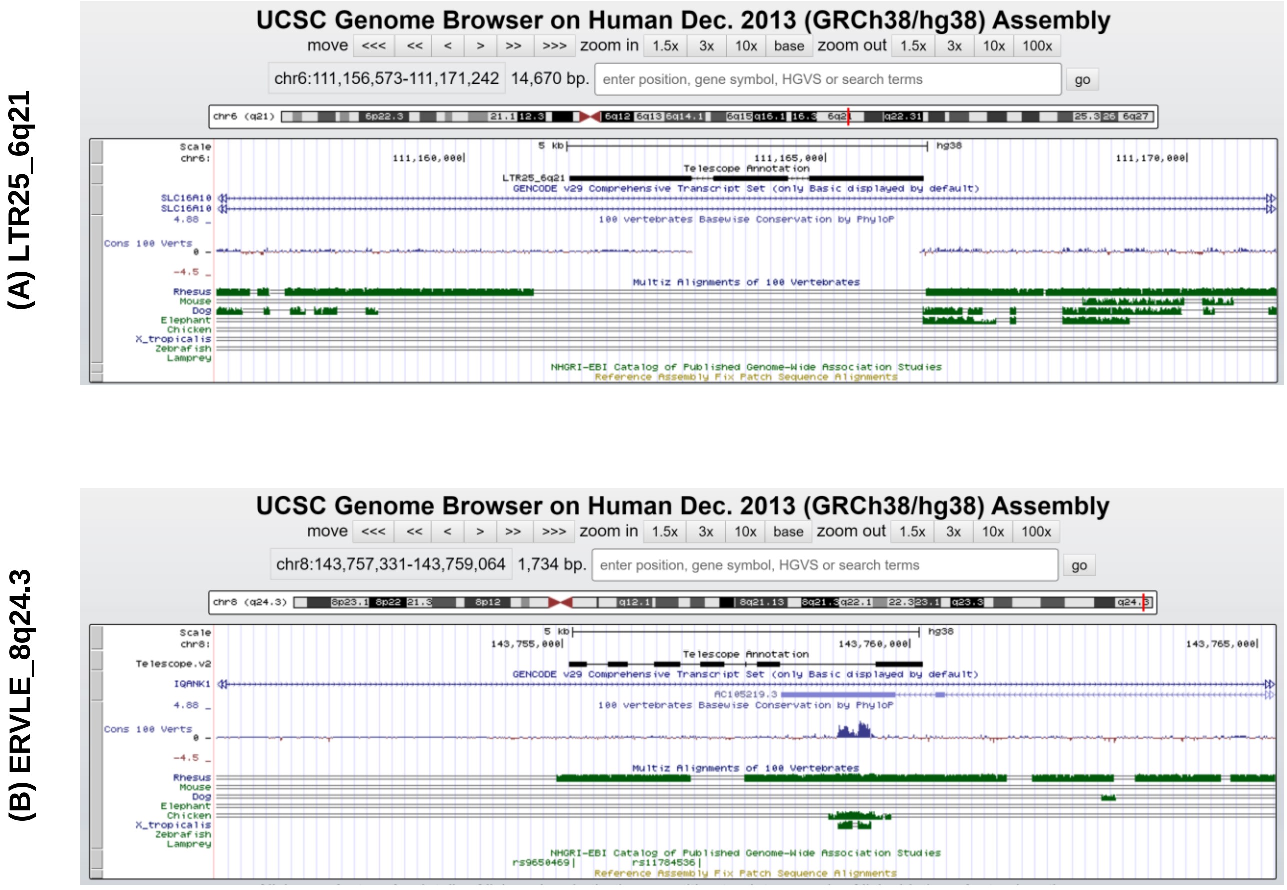


**Supplemental Figure 6.** Genomic context of the two HERVs implicated in schizophrenia, as observed in the UCSC browser. **(A)** LTR25_6q21 is in an intronic region of a gene that encodes a protein responsible for transportation of aromatic amino acids across the plasma membrane (*SLC16A10*), although is encoded in the antisense strand. **(B)** ERVLE_8q24.3h is in an intronic region of a gene that encodes a protein responsible for (*IQANK1*), although is in the antisense strand. The “Cons 100 Verts” track corresponds to sequence conservation across 100 vertebrates, and the tracks below show conservation across individual organisms, including rhesus macaque, mouse, dog, elephant, chicken, frog (Xenopus tropicalis), zebrafish and lamprey. These data suggest that LTR25_6q21 is exclusive to humans, and ERVLE_8q24.3h evolutionarily conserved in primates.

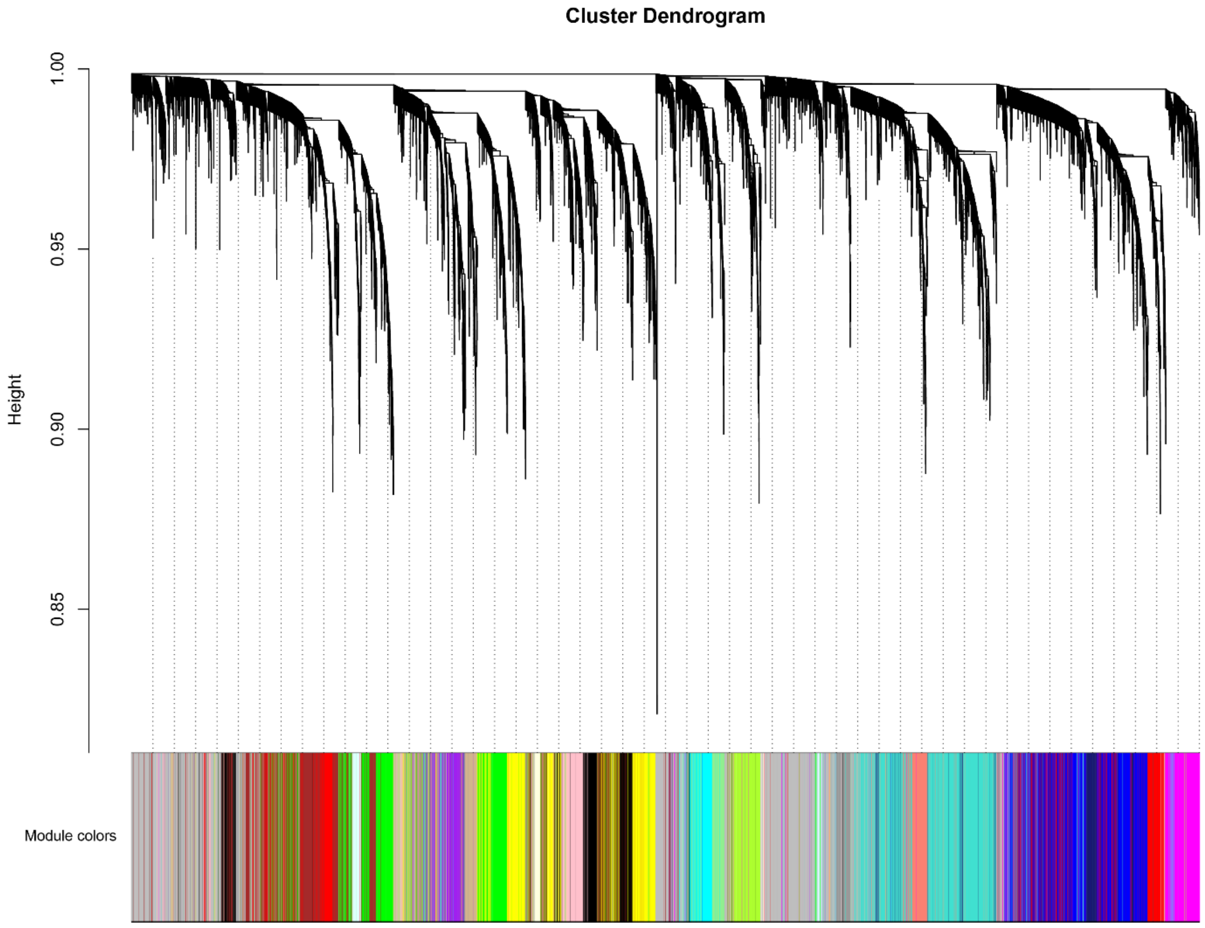


**Supplemental Figure 7.** Clustering dendrogram of genes and HERVs expressed in the human DLPFC (n = 279 controls, n = 259 schizophrenia patients), with dissimilarity based on topological overlap, and the assigned module colors below. HERV and gene counts were variance stabilized transformed in DESeq2 [5] and controlled for the effect of all main confounders known to affect gene expression in the brain using the limma [6] package (see Methods for further details).

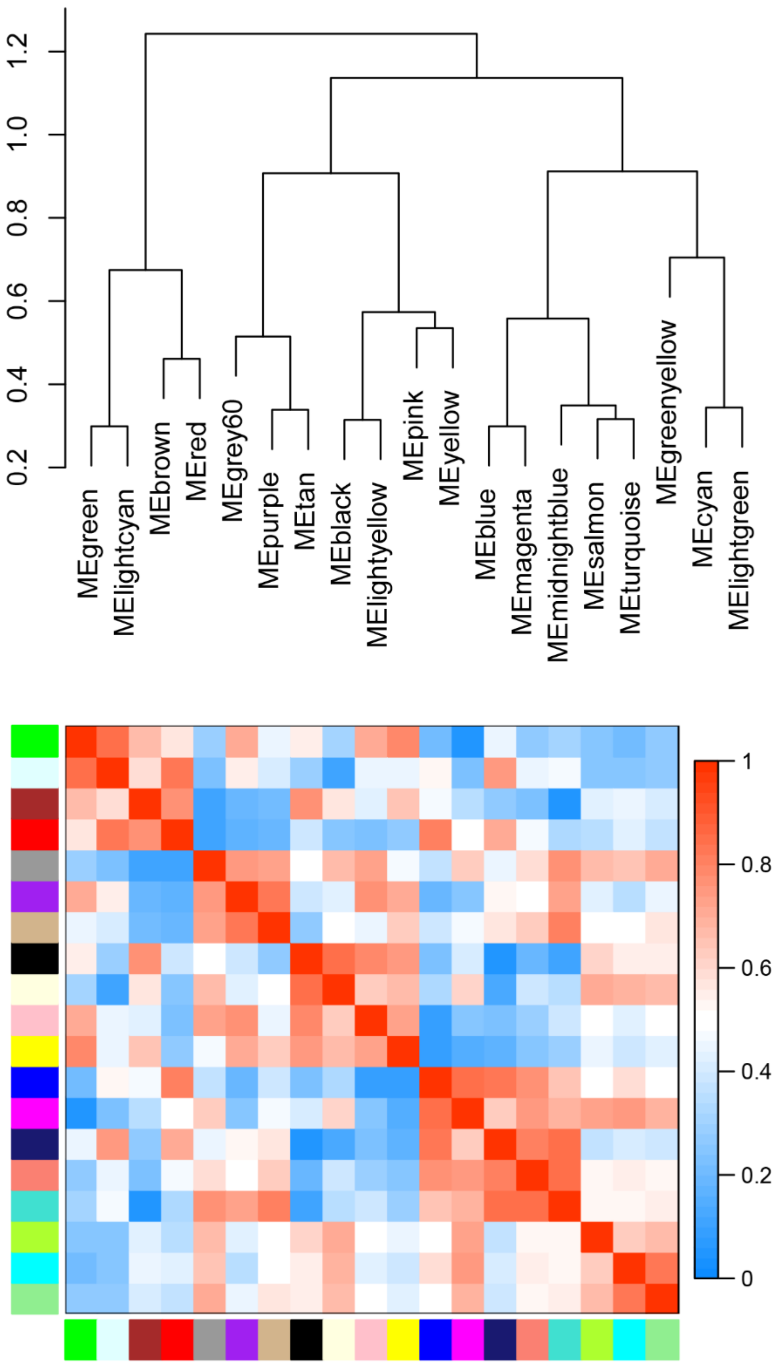


**Supplemental Figure 8.** Relationship amongst the co-expression modules detected in the DLPFC samples (n = 279 controls, n = 259 schizophrenia patients). **(A)** Hierarchical clustering dendrogram of the eigengenes according to module adjacency. **(B)** Eigengenes adjacency represented as a heatmap. The function of closely related modules is likely to be related. For example, the green module (where LTR25_6q21 belongs) is positively correlated with the red module, and both modules are enriched for gene ontology terms associated with synapse and neuronal function.

### ***Supplemental Methods***

#### Brain samples

Human dorsolateral prefrontal cortex (DLPFC) sections (Brodmann Area 9) were obtained from the Medical Research Council London Neurodegenerative Diseases Brain Bank at the Institute of Psychiatry, Psychology & Neuroscience, King's College London, under license issued by the United Kingdom Human Tissue Authority (ref. 12293). This cohort consisted of 8 females and 2 males; the mean age of the subjects was 68 (range: 55-77 years); the period of post-mortem delay per subject consisted of 52 hours on average (range: 21-78hrs).

#### Cell culture

We used the neural stem cell line CTX0E16 to investigate HERV expression during in vitro differentiation. This cell line was kindly provided by ReNeuron (www.reneuron.com) as part of a Material Transfer Agreement. This cell line was derived from the brain of a 12-week male fetus as detailed elsewhere [4,7]. Cells were routinely maintained at the proliferative stage, or terminally differentiated into neurons for 28 days (DD28), as described elsewhere [2-4]. Cultures were kept at 37 $^{o}$C, with 5% $CO_2$, and split using Accutase when necessary. The proliferative state was maintained by culturing cells in DMEM:F12 (Sigma) supplemented with 0.03% human serum albumin (PAA), 100 $\mu g.ml^{-1}$ apo-transferrin (Scipac), 16.2 $\mu g.ml^{-1}$ putrescine (Sigma), 5 $\mu g.ml^{-1}$ human insulin (Sigma), 60 $ng.ml^{-1}$ progesterone (Sigma), 2 mM L-glutamine (Sigma) and 40 $ng.ml^{-1}$ sodium selenite (Sigma), 10 $ng.ml^{-1}$ human FGF2, 20 $ng.ml^{-1}$ human EGF (PeproTech) and 100 nM 4-hydroxy-tamoxifen (4-OHT) (Sigma). Terminal differentiation of these cells was achieved by replacing DMEM:F12 medium with Neurobasal Medium supplemented with B27 serum-free supplement (Life Technologies), 0.03% human serum albumin, 100 $\mu g.ml^{-1}$ apo-transferrin, 16.2 $\mu g.ml^{-1}$ putrescine, 5 $\mu g.ml^{-1}$ human insulin, 60 $ng.ml^{-1}$ progesterone, 2 mM L-glutamine, and 40 $ng.ml^{-1}$ sodium selenite. Neural progenitor cells were grown on laminin-coated (1 $\mu g/cm^2$) T75 Nunc flasks, and differentiated cultures were grown on flasks coated with poly-D-lysine (0.2 $mg.ml^{-1}$, Sigma) and laminin.

***RNA isolation and cDNA synthesis***

Total RNA was extracted using Tri-Reagent (Thermo Fisher Scientific), according to the manufacturer's protocol. An additional step of sample homogenization in this reagent was applied for extraction of RNA from brain samples, which used the Omni Polytron homogenizer (G5-95 5mm probe) coupled to a PowerGen 125 rotor (single 45 sec step). RNA samples were analysed in a spectrophotometer NanoDrop ND-1000 (Thermo Fisher Scientific), and in a 2100 Bioanalyzer (Agilent; Santa Clara, California, USA) using the Agilent RNA 6000 Pico Kit. On average, brain samples showed absorbance ratios of 1.9 and RNA Integrity Numbers (RIN) of 4.9, whereas samples from tissue culture were associated with RIN of 10.0. Total RNA samples were treated using the TURBO DNA-free kit (Thermo Fisher Scientific) and the DNase Inactivation Reagent, according to the manufacturer's protocol. This RNA did not yield a PCR product for beta-actin in the absence of a reverse transcription step, confirming removal of contaminating DNA. Reverse transcription was performed using 1.0-2.0 µg of DNA-free RNA and SuperScript III Reverse Transcriptase and RNaseOUT Recombinant Ribonuclease Inhibitor (Thermo Fisher Scientific), according to the manufacturer's protocol. Samples were stored at -80ºC until use.

***Quantitative reverse transcription polymerase chain reaction (RT-qPCR)***

HERV expression was measured in a QuantStudio 7 Flex or a QuantStudio 5 Real-Time PCR system (Thermo Fisher Scientific) using the relative standard curve method in the QuantStudio Real-Time PCR Software v1.3 or the QuantStudio Design & Analysis Software, respectively. The reactions were run in triplicates, and consisted of 24 ng cDNA, HOT FIREPol EvaGreen qPCR Mix Plus (Solis BioDyne, Tartu, Estonia) and 200 nM primers (final volume = 12.5 µl reactions). The thermocycling conditions consisted of a 15 min incubation step at 95 ºC, followed by 40 cycles of 30 sec at 95 ºC, 30 sec at 62.5 ºC, 30 sec at 72 ºC, followed by a melting curve analysis using a continuous temperature gradient from 60 ºC to 95 ºC. Melting curves revealed single, clear peaks for each primer set (**Supplemental Figure 2C, Supplemental Figure 5C**), and PCR efficiencies were all between 1.8 and 2.1, as determined by standard curves of pooled cDNA using five 1:2 dilution points. Primers were designed using Primer3 [8] and purchased from Integrated DNA Technologies (Leuven, Belgium) (**Supplemental Table 11**). To generate HERV primers with single locus resolution, we generated family alignments using Geneious v4.8.5 (Biomatters Ltd, Auckland, New Zealand) and ClustalW2, so that primers could be designed within fragments that were exclusive to the HERV locus of interest. Expression was normalized to a standard curve of five 1:2 dilution points, and is relative to the geometric mean of the normalized expression of the housekeeping genes analyzed in each experiment (*ACTB*, *SDHA* and *ALG2*, in the adult brain analysis; and *RPL6* and *ALG2* in the cell line analysis).

### *References*

1. Fromer, M. *et al.* Gene expression elucidates functional impact of polygenic risk for schizophrenia. *Nature neuroscience* **19**, 1442-1453 (2016).
2. Deans, P.J.M. *et al.* Psychosis Risk Candidate ZNF804A Localizes to Synapses and Regulates Neurite Formation and Dendritic Spine Structure. *Biol Psychiatry* **82**, 49-61 (2017).
3. Duarte, R.R.R. *et al.* The Psychiatric Risk Gene NT5C2 Regulates Adenosine Monophosphate-Activated Protein Kinase Signaling and Protein Translation in Human Neural Progenitor Cells. *Biol Psychiatry* **86**, 120-130 (2019).
4. Anderson, G.W. *et al.* Characterisation of neurons derived from a cortical human neural stem cell line CTX0E16. *Stem Cell Res Ther* **6**, 149 (2015).
5. Zhu, A., Ibrahim, J.G. & Love, M.I. Heavy-tailed prior distributions for sequence count data: removing the noise and preserving large differences. (2018).
6. Ritchie, M.E. *et al.* limma powers differential expression analyses for RNA-sequencing and microarray studies. *Nucleic Acids Research* **43**, e47-e47 (2015).
7. Pollock, K. *et al.* A conditionally immortal clonal stem cell line from human cortical neuroepithelium for the treatment of ischemic stroke. *Exp Neurol* **199**, 143-55 (2006).
8. Untergasser, A. *et al.* Primer3--new capabilities and interfaces. *Nucleic Acids Res* **40**, e115 (2012).
